# Synfire Chain Dynamics Unravelling Theta-nested Gamma Oscillations for Balancing Prediction and Dodge in Navigation

**DOI:** 10.1101/2024.03.01.583075

**Authors:** Kwan Tung Li, Yina Wei, Pulin Gong, Dongping Yang

## Abstract

Theta-nested gamma oscillations, widely observed in experiments, play a crucial role in navigation, yet their functional roles and the origin of the positive correlation between theta frequency and motion velocity remain unclear. We propose that the object’s survival relies on both prediction and dodge – predicting future events and staying alert to unpredictable ones, the latter of which has seldom been considered in goal-navigation tasks. By building a biologically plausible spiking neuronal network model and reproducing experimental results, we leverage synfire chain properties – length and separation – to elucidate the functional roles of theta-nested gamma oscillations: theta oscillations for self-location awareness, gamma oscillations for predictive capabilities and their coupling for enhancing functionality. The positive correlation between theta frequency and motion velocity is demonstrated to optimally balance representing predictable events for planning and staying alert to unexpected events. Our study offers a new avenue for unravelling the neural mechanisms of navigation.

## 1 Introduction

During navigation, theta-nested gamma oscillations, characterized by faster gamma oscillations nested within slower theta oscillations, are frequently observed in neural activities [1–3]. Impairment of theta or gamma oscillation has been linked to deficits in navigation performance [4–6]. However, the functional roles of theta and gamma oscillations and their coupling during navigation remain ambiguous: Some studies suggest that theta-phase gamma-amplitude coupling correlates with memory performance [7–9], with the theta period playing a role in chunking spatial information [7, 8], while others propose that theta and gamma oscillations synchronize networks over long and short distances, respectively [10, 11]. An even more perplexing finding in experiments is that theta frequency in theta-nested gamma oscillations positively correlates with motion velocity during navigation [12–16]. Whether such a correlation can be intrinsically related to navigation performance remains unexplored.

Here, we propose that the functional roles of theta-nested gamma oscillations during navigation can be disclosed, if one considers the ability to predict future events, e.g., navigating towards rewarding locations like those offering food or shelter, and the need to remain alert to the surrounding environment are equally significant in goal navigation. Predicting the future as an essential survival skill relies on the brain’s ability to remember the location of these reward sites and the corresponding paths [1, 17], yet the journey to these rewards can be fraught with *unpredictable events* such as predators, traps, and accidents. Nevertheless, the unpredictable events were overlooked in previous studies [3, 17, 18]. This particular perspective provides a critical starting point for uncovering the functional roles of theta-nested gamma oscillations in navigation.

To achieve this, we conducted computational simulations of goal navigation with both predictable and unpredictable events. We introduced a biologically plausible spiking neural network model, where synfire chains [19, 20] were instrumental in unravelling the impact of theta-nested gamma oscillations on navigation performance. Delving into the dynamics of synfire chains, we examined varying theta frequencies and motion velocities, exploring normal conditions and the consequences of impaired theta or gamma oscillations. Our investigation highlights the inherent capacity of synfire chains to anticipate future events (*synfire chain length*) and emphasizes the need for heightened awareness of surroundings (*synfire chain separation*). Our results elucidate that theta oscillations contribute to spatial information chunking and self-location recognition, fostering alertness, while gamma oscillations facilitate synfire chain propagation over short distances, synchronizing neural activity to enhance predictive abilities. Importantly, our findings unveil a self-organized mechanism wherein the positive correlation between theta frequency and motion velocity serves as an optimal trade-off between predicting future events and remaining alert to the surroundings. Our study offers a new perspective for unravelling the functional role of theta-nested gamma oscillations in navigation, by considering both predictable rewards and unpredictable challenges.

## 2 Results

### 2.1 Sketch of Our Biologically Plausible Network Model

During navigation, the interplay of theta-nested gamma oscillations orchestrates synfire chains, which emerge within theta cycles, with discrete clusters of place cells activating across gamma cycles and each cluster encoding a specific location. Our investigation commenced by building a biologically plausible network model to reproduce synfire chain properties, which is a sequential firing with direction, and theta-nested gamma oscillations (Fig. 1).

**Fig. 1.**
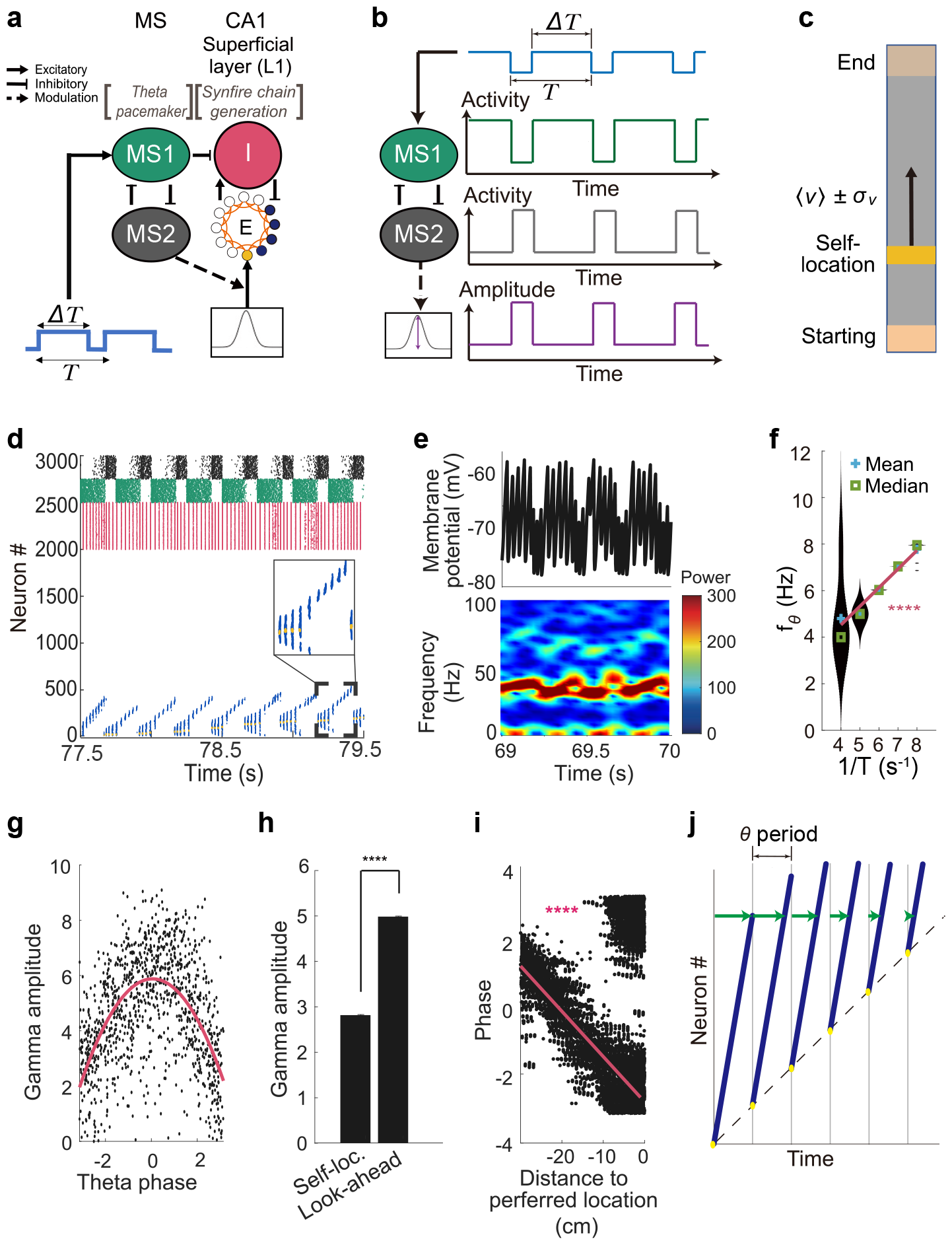
Synfire chain dynamics generating theta-nested gamma oscillations. (a) Schematic depiction of the model architecture. (b) MS modulating location-dependent input. (c) Linear track. (d) Spike raster plot of 2000 excitatory neurons representing self-location (yellow) and prediction (blue), 500 inhibitory neurons (red) in L1, and 250 inhibitory neurons in both MS1 (green) and MS2 (black). (Insert): Manification. (e) Population mean membrane potential of excitatory neurons (Top) and corresponding power spectrum (Bottom). (f) Linear dependence of *f*_*θ*_ in L1 on *T* . The width of the violin plot is normalized to an identical scale. (g) Coupling of theta phase and gamma amplitude. (h) Gamma amplitudes at self-location and look-ahead part of synfire chains. (i) Theta-phase precession. (j) Illustration of phase precession. Dot line: motion trajectory; Yellow dots: self-location of synfire chains; Blue lines: look-ahead part of synfire chains; The separation between consecutive gray lines represents one theta period; The green arrow length denotes the phase displacement. Parameter used: *v* = 10 cm/s and *T* = 250 ms. Data in (f, h) are represented as mean *±* standard error of the mean (SEM). *n* = 10 trails/setting. ****: *p <* 0.0001. In (f), two-sample Student’s t-test; In (i), Pearson’s correlation.

This model encompasses two crucial components: the medial septum layer (MS) and the CA1 superficial layer (L1) (see Fig. 1a). MS generates theta oscillations, while L1 is responsible for synfire chain formation. Within MS, two interconnected inhibitory populations, MS1 and MS2 [21–23], play key roles. MS1 received tonic inputs—potentially originating from an excitatory-inhibitory (E-I) circuit within the MS [21]—to regulate theta frequency, which is modelled as a step function with period *T* including up state period Δ*T* and down state period *T*− Δ*T* (Fig. 1a). In L1, an E-I network operated, with excitatory neurons receiving location-dependent inputs from the entorhinal cortex (EC) [24]. These MS2-modulating inputs [25] were modeled as Gaussian profiles across the spatial domain, and exhibiting an out-of-phase relationship with CA1 theta oscillations [26, 27] (Fig. 1a, b). The next subsection investigated synfire chain dynamics by simulating a rat navigating a linear track at a velocity *v* with its mean ⟨*v*⟩ and standard deviation *σ*_*v*_ (Fig. 1c). Accordingly, the excitatory network in L1 was configured as a continuous attractor neural network (CANN) (Fig. 1a). When MS1 was activated by the up state of the step function, MS2 and inhibitory population in L1 were suppressed. The suppressed MS2 and inhibitory population weakened the MS2-modulating inputs and provided high excitability in L1 excitatory population respectively, initiating the synfire chain propagation. (Fig. 1b, d). On the other hand, when MS1 was at suppressed, MS2 and inhibitory population in L1 were at high activity. The high inhibition from inhibitory population reduced the excitability of excitatory neurons in L1, while the high activity of MS2 strengthened the MS2-modulating inputs. As a result, the excitatory neurons were driven by the location-dependent input and fired around the current location of the rat (Fig. 1b, d). Further methodological details are provided in Sec. 4.

From subsection 2.3 onward, our investigation delved into the nuanced dynamics of goal-navigation tasks within a cross-maze paradigm (Fig. 2). The tasks involved finding a reward (*a predictable event*) and avoiding a trap (*an unpredictable event*), as depicted in Fig. 2a. The maze comprised four sections: the main track, the left, middle and right arm, each spanning 50 cm. The reward was placed within one of the arms, while the trap was randomly positioned along the main track.

**Fig. 2.**
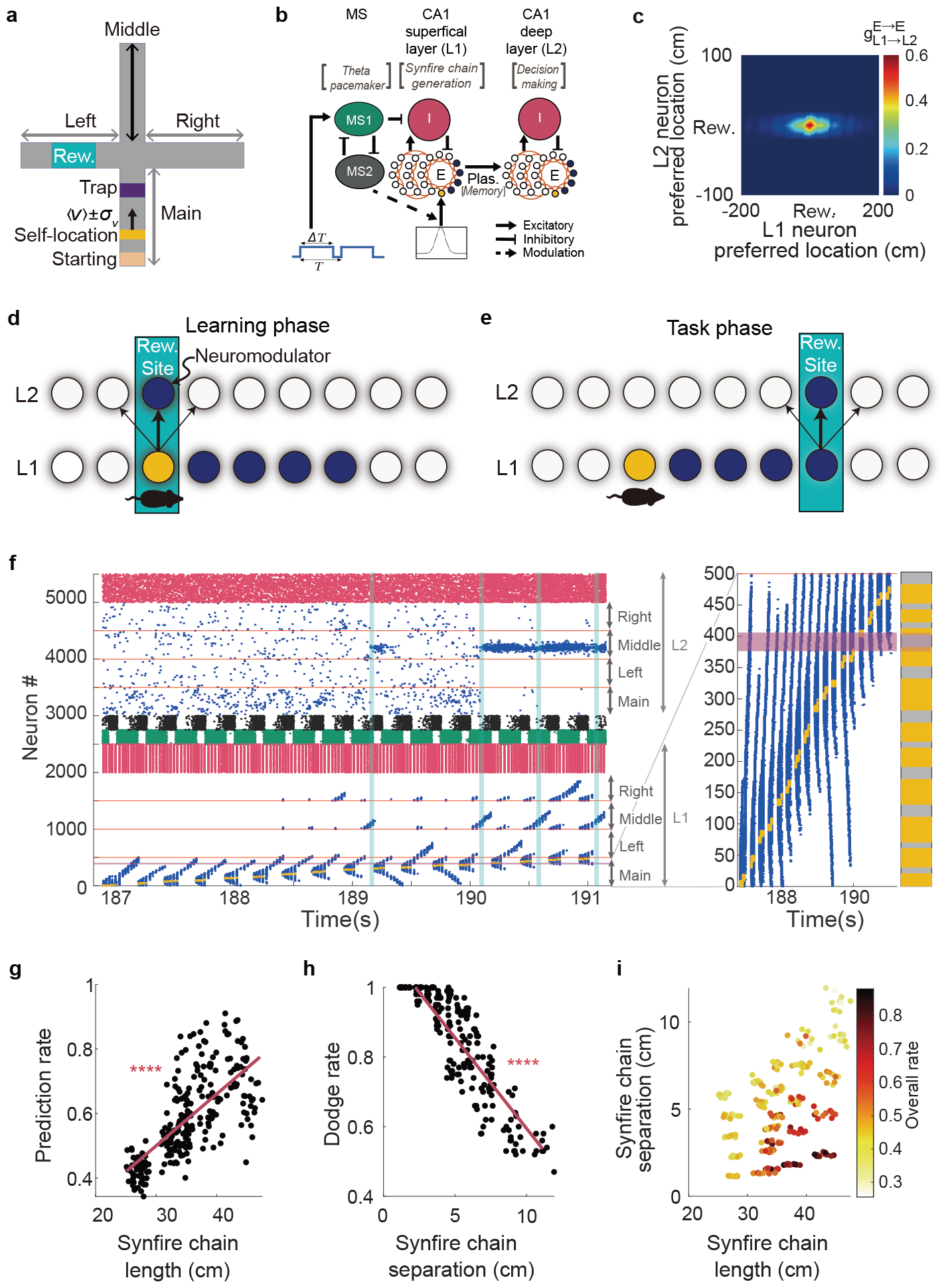
Goal-navigation task, synfire chain properties and navigation performance. (a) Cross maze with a main track and three batches; Each section is 50 cm long with periodic boundary conditions: Terminations of the three batches are connected to the starting point of the main track. (b) Schematic depiction of the adapted model architecture with adding L2: Excitatory circuits in L1 and L2 consist of three CANNs, each representing a distinct trajectory: from the main track to the left, the middle and the right, respectively; They collectively share the neurons representing the main track, representing diverse navigational paths. (c) Synaptic weights from excitatory neurons of L1 to those of L2 after the learning phase. (d, e) Illustration of successful learning during the learning phase (d) and prediction during the task phase (e). rat symbol: current location of the rat; Yellow: self-location of synfire chain; Blue: look-ahead part of synfire chain; The thickness of the arrow between L1 and L2 represents the synaptic weight. (f) Spike raster plot with the same color notation as in Fig. 1d. Cyan bars: The reward site in L2 is being activated by a synfire chain in L1 (Left). Magnification of the main track and projecting all the self-location and trap at the far right (Right). (g-i) Effects of synfire chain properties on goal-navigation performance at various *v* and *f*_*θ*_: Effects of synfire chain length on prediction ability (g); Effects of synfire chain separation on dodge ability (h); Their combined effects on the overall probability of reaching reward (i). ****:*p <* 0.0001. In (g, h), Pearson’s correlation.

To explore the efficacy of synfire chains in goal-navigation tasks, we extended our model by incorporating a deep layer (L2) into CA1 (Fig. 2b). L2 functioned to store information about reward locations [28], facilitating decision-making regarding which arm to navigate toward at the maze’s fork. L2 comprised an E-I circuit that received plastic inputs from L1. Consequently, excitatory circuits in both L1 and L2 were adapted to include three CANNs, each representing a distinct trajectory: from the main track to the left, middle or right arm, respectively; These networks representing various navigational paths shared the neurons associated with the positions in the main track. For further information, refer to Sec. 4.

### 2.2 Synfire Chain Dynamics Generating Theta-nested Gamma Oscillations

Within the cognitive map, a conceptual framework representing the subject’s environment [29–32], the synfire chain assumes a critical role in efficiently recalling and predicting navigation trajectories, contributing to the retrieval and utilization of spatial information [1–3, 17]. It enables rapid traversal through the cognitive map, facilitating the anticipation and execution of movements and decisions [3, 18].

In our model, synfire chains can be divided into two parts: One encodes the subject’s current location (self-location, yellow dots in Fig. 1d), and the other predicts future locations (look-ahead locations, blue dots in Fig. 1d), predominantly advancing forwards. Synfire chains were initiated, terminated and re-initiated at the subject’s self-location by activating MS1. This activation inhibited MS2 and L1 inhibitory neurons, inducing a period of heightened excitability within L1 excitatory neurons, thereby facilitating synfire chain generation. In L1, excitatory neurons exhibited higher connectivity to nearby place cells (Sec. 4), wherein initially activated place cells underwent adaptation to decrease their activities and enhanced the excitability of adjacent place cells, rendering the propagation of excitation.

The dynamics of synfire chains exhibited theta-nested gamma oscillations (Fig. 1e), with the theta rhythm originating from MS and gamma oscillations from the E-I loop in L1 [33, 34]. The tonic inputs received by MS1, temporally separated by *T* (Fig. 1a, b), generated synfire chains in L1 with various theta frequencies *f*_*θ*_, nearly equal to 1*/T* (Fig. 1f). The amplitude of gamma oscillations was modulated by the theta phase (Fig. 1g), consistent with experimental findings [9]. The modulation resulted in a larger gamma amplitude during the look-ahead phase compared to the self-location phase (Fig. 1h), as deactivation of MS1 during the latter increased inhibition on excitatory neurons in L1, thus suppressing membrane potential oscillations.

Furthermore, our simulation revealed the occurrence of theta-phase precession (Fig. 1i), wherein a place cell gradually shifts its firing phase to an earlier phase within the subsequent theta cycle [2, 3, 35]. In our model, theta-phase precession was induced by consecutive synfire chains originating from locations near the place field of the respective place cell [2]. The synfire chain encoded successive locations, from the subject’s current position to future locations. As the subject advanced, the same place cell approaches its preferred location in the forthcoming theta cycle, leading to activation at an earlier phase during forward movement (Fig. 1j).

### 2.3 Learning the Association of Reward Site between L1 and L2

Simulating goal-navigation tasks in a cross maze involved two distinct phases: the learning phase and the task phase.

During the learning phase, a simulated rat, representing the subject, explored the maze. Upon reaching the reward site, neuromodulator signals targeted corresponding excitatory neurons in L2 (Fig. 2c, d), initiating reward spike-timing dependent plasticity (R-STDP) [36–38]. The learning mechanism strengthened weak synapses but weakened strong ones (Supplementary Fig. 1a), when subjected to neuromodulator signals and pre-post spike correlation (Sec. 4). Specifically, synapses connecting L1 and L2 at the reward site underwent potentiation (Fig. 2c) under the combined influence of the neuromodulator and pre-post spikes (Supplementary Fig. 1b, d). Conversely, synapses between neurons neighboring the reward site remained unaffected due to the absence of neuromodulators (Supplementary Fig. 1c, e). Consequently, neurons in L2 displayed heightened in response to the L1 input around the reward site, consistent with the idea that L2 is predominantly involved in processing reward-related information [28].

In the subsequent task phase, the rat revisited the maze without R-STDP. Upon reaching the reward location, the synfire chain in L1 triggered a high firing rate in L2 around the reward site via the strengthened connections from L1 (Fig. 2e, f, cyan bars), which had been acquired during successful learning (Fig. 2d). This resulted in a notable disparity in population firing rates between the rewarded arm and the other arms in L2, thereby elevating the probability of the rat selecting the arm with the reward (*prediction rate*). Conversely, if the synapses between L1 and L2 were not potentiated, the firing rate among the three arms would be similar, resulting in a low prediction rate.

Upon the rat approaching the fork, a synfire chain explored one of the arms. The probability of selecting a specific arm was determined by the firing rate of L2 neurons representing that arm. Meanwhile, the corresponding place cell experienced adaptation (Sec. 4), decreasing the probability of re-entering the same arm in subsequent synfire chains, enabling exploration of different arms (Fig. 2f). This phenomenon aligns with vicarious trial and error (VTE) observed in experimental settings [3, 18]. An imperceptible trap was randomly positioned along the main track to simulate unpredictable events, becoming perceptible to the rat as it approached closely. If the synfire chain’s self-locations did not coincide with the trap location when the rat passed this location, it became trapped, impeding its progress towards the reward. Otherwise, it would evade the trap and continue its quest for the reward (*dodge rate*).

### 2.4 Goal-navigation Performance Determined by Synfire Chain Properties

The goal-navigation performance can be evaluated by the overall success rate of reaching the reward, represented by the combined efficacy of the prediction and dodge rates. This evaluation enables the study of how theta frequencies (*f*_*θ*_) and motion velocities (*v*) affected the performance. Various spike raster plots were illustrated in Supplementary Fig. 2. Synfire chains also exhibited distinct characteristics at different *v* and corresponding *f*_*θ*_: Higher velocity led to shorter synfire chains (*synfire chain length*) and closer spatial intervals between consecutive synfire chains (*synfire chain separation*) (Supplementary Fig. 2).

Significantly, synfire chain properties were closely associated with the performance: A positive correlation was observed between synfire chain length and the ability to predict future events (Fig. 2g), and a negative correlation existed between synfire chain separation and the dodge ability (Fig. 2h). Intuitively, longer chains extended further towards the reward, thereby enhancing the prediction ability, and larger separations diminished the ratio between self-location and look-ahead, leading to a decrease in the dodge ability. Therefore, optimizing the overall rate of reaching rewards crucially hinges on achieving longer synfire chains with closer separations (Fig. 2i).

Building upon this framework, we investigated the functional significance of theta and gamma oscillations in the goal-navigation performance in the following two subsections. These investigations were conducted under the assumption of a positive correlation between *f*_*θ*_ and *v*, which was widely observed in experiments [12–16] and to be demonstrated for optimally balancing prediction and dodge in Sec. 2.7.

### 2.5 Theta Impairment Inducing Loss of Self-location

Experimentally, theta oscillation impairment can be achieved by injecting muscimol, a GABA receptor agonist, to inactivate MS in rats [4, 6]. This resulted in reduced theta rhythm, deteriorated navigation performance, and unrelated firing of place cells with the rat’s actual location. The underlying mechanism can be elucidated by conducting simulations with augmenting the synaptic conductance between MS1 and MS2 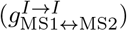, as well as within MS1 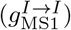 and MS2 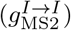, by a factor 3.

This manipulation indeed resulted in a noticeable decrease in overall activity within MS, particularly in MS2 (Supplementary Fig. 3a, b). The prolonged activation of MS1 and low MS2 activity enhanced the look-ahead part of synfire chains (Fig. 3b), as it hindered the ability of MS2 to terminate the synfire chain, resulting in a prolonged unitary synfire chain. This disruption caused the self-location to no longer align with the beginning of the chain (Fig. 3a, b), even though the periodic boundary condition induced multiple coincident self-location phases, a phenomenon absent in an infinite maze.

**Fig. 3.**
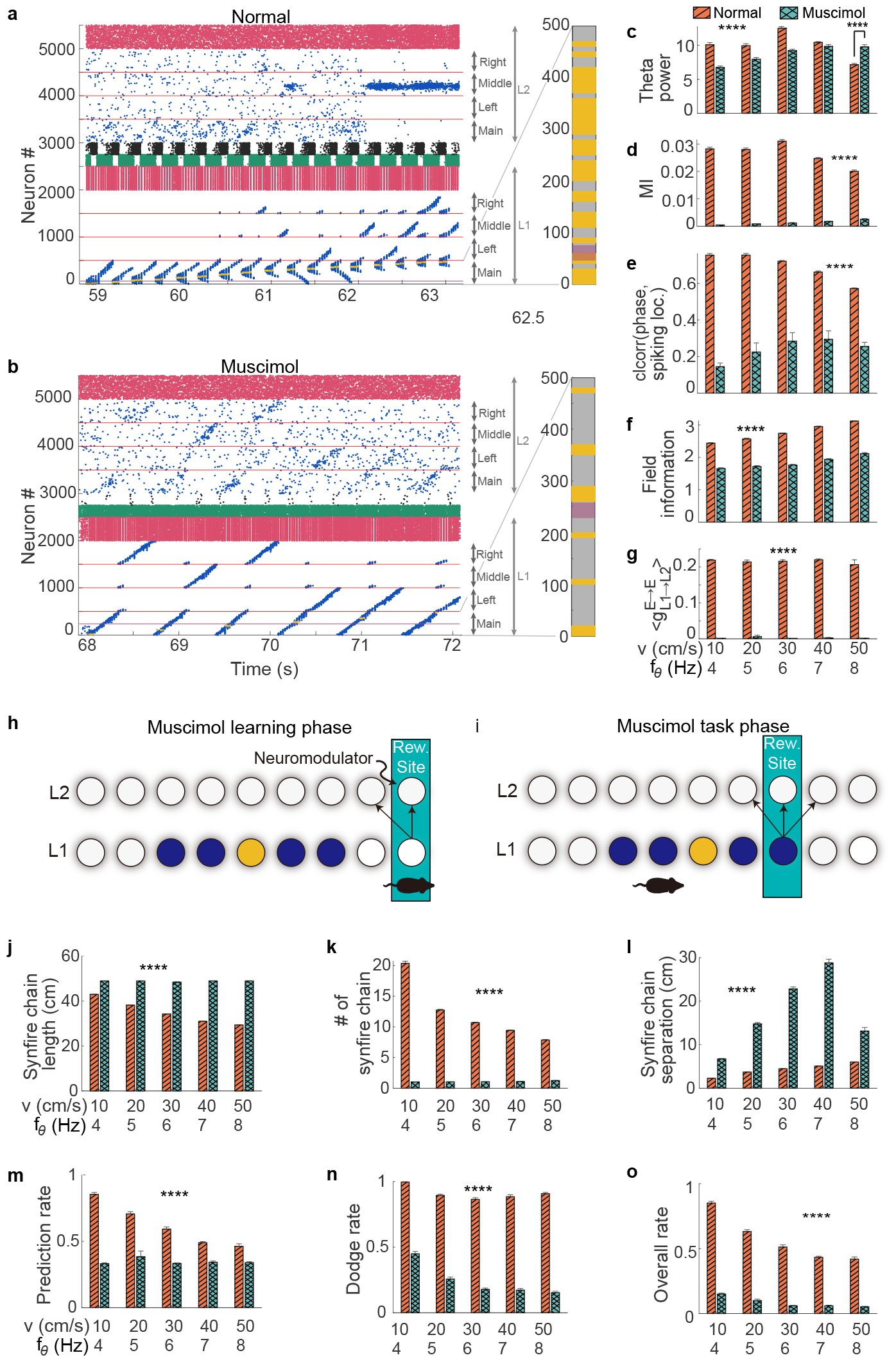
Theta oscillation impairment. (a, b) Raster plots (Left) and projection of self-location (right) of Normal (a) and Muscimol condition (b) at *v* = 10*cm/s* and *f*_*θ*_ = 4*Hz*, with the same color notation as that in Fig. 1d. (c-g, j-o) Comparisons between normal and muscimol conditions: (c) Theta power; (d) Modulation index (MI); (e) Circular-linear correlation (clcorr) between theta phase and spiking location; (f) Field information; (g) Averaged excitatory to excitatory synaptic weight from L1 to L2 at the reward site. (h, i) Illustration of learning phase (h) and task phase (i) when muscimol is applied. Notifications are the same as those in Fig1.12d. (j) Synfire chain length. (k) number of synfire chains between the starting point and the fork. (l) Synfire chain separation between consecutive synfire chains. (m) Prediction rate. (n) Dodge rate. (o) The overall rate of receiving the reward. Data in (c-g, j-o) are represented as mean *±* SEM. *n* = 10 trails/setting. ****: *p <* 0.0001. In (d-g, j-o) and *v* = 10 ∼ 40 cm/s in (c), 2ANOVA test; *v* = 50 cm/s in (c), Two-sample Student’s t-test.

Meanwhile, the reduced activity of MS2 weakened its modulation on location-specific inputs and diminished the activity of inhibitory neurons in L1, leading to reduced theta power, except during fast movement (Fig. 3c). Quantitative analyses (Sec. 4) employing modulation index (MI) and circular-linear correlation (clcorr) indicated muscimol injection decreased theta-phase-gamma-amplitude coupling, and less significant theta-phase precession (Fig. 3d, e). Furthermore, muscimol application reduced field information (Fig. 3f), signifying a decreased correlation between spike activity and the subject’s current location. These results were consistent with experimental findings [3–5, 39].

During the learning phase, synapses from L1 to L2 at the reward site had not been potentiated (Fig. 3g), as the disrupted synfire chain impaired spike emission in L1 upon neuromodulator release at the reward site and rendered neurons in L2 unresponsive to synfire chains initiated in L1 (Fig. 3h, i). During the subsequent task phase, the disrupted synfire chain was unable to activate neurons in L2 at the reward site (Fig. 3b, i), leading to similar firing rates among arms and a correspondingly lower prediction rate (Fig. 3m), consistent with experimental results [4]. Notably, the administration of muscimol induced longer synfire chains (Fig. 3j) but with a significant reduction in their total number before reaching the fork (Fig. 3k), impairing learning during navigation (Fig. 3g, h). Thus, the inhibited MS led to a lower dodge rate (Fig. 3n) due to increased synfire chain separation (Fig. 3l), ultimately diminishing the overall rate of reward acquisition (Fig. 3o). These findings suggest that inhibiting MS disrupted the synfire chain’s ability to memorize the reward site location, impeding the goal-navigation performance.

### 2.6 Gamma Impairment Inducing Weak Synchronization and Theta-gamma Coupling

Impairment of gamma oscillation has been hypothesized to degrade the quality of theta-phase precession [5]. The impact of gamma oscillations [33, 40] was investigated here by increasing the excitatory synaptic decay time constant 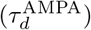 from 5 ms (Normal) to 20 ms (NO gamma) (Fig. 4a, b), resulting in reduced gamma power (Fig. 4c) and synchronization (Supplementary Fig. 4a), accompanied by a decrease in modulation index (MI) (Fig. 4d), and a decline of circular-linear correlation between theta phase and spiking location (Supplementary Fig. 4b). These results indicated a decrease in theta-gamma oscillation coupling and theta-phase precession, consistent with the hypothesis [5].

**Fig. 4.**
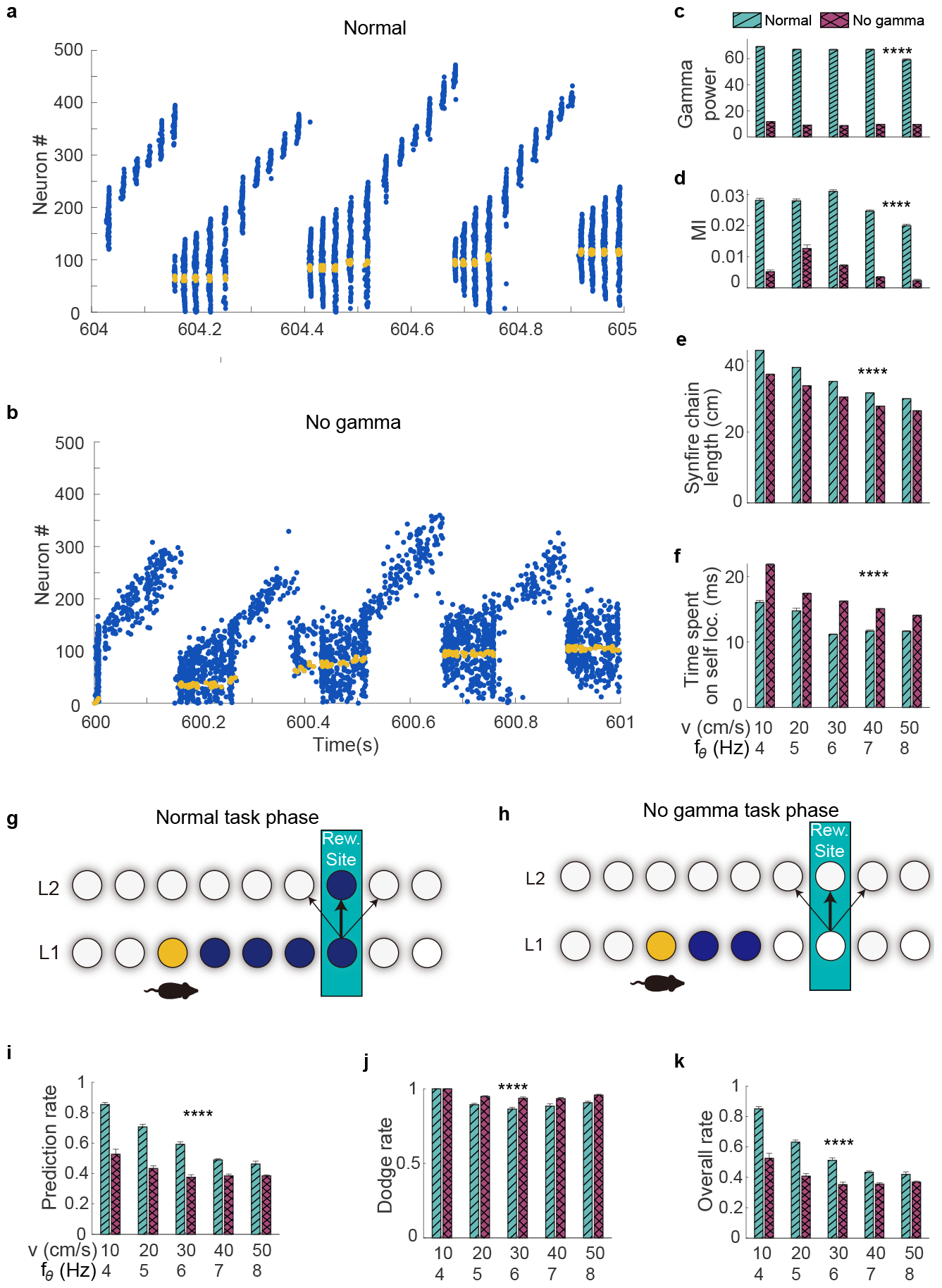
Gamma oscillation impairment. (a, b) Raster plots of Normal (a) and No gamma (b) at *v* = 10 cm/s and *f*_*θ*_ = 4 Hz, with the same color notation as that in Fig. 1d. Normal: 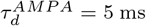; No gamma: 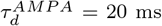. (c-f, i-k) Comparisons between Normal and No gamma: (c) Gamma power; (d) Modulation index (MI); (e) Synfire chain length; (f) Time spent on self-location, indicated by the width of yellow dots in (a, b). (g, h) Illustrations of task phase at Normal (g) and No gamma (h). Notifications are the same as those in Fig. 2d. (i) Prediction rate. (j) Dodge rate. (k) The overall rate of receiving rewards. Data in (c-e, i-k) are represented as mean *±* SEM. *n* = 10 trails/setting. ****: *p <* 0.0001. In (c-e, i-k), 2ANOVA test.

In our simulation, NO gamma led to shorter synfire chains (Fig. 4e), with a minor effect on learning (Supplementary Fig. 4c), and minor changes in the number of synfire chains between the starting point and the fork (Supplementary Fig. 4d). Notably, the response time to the removal of the stimulus was prolonged, resulting in a temporal extension of the self-location part of synfire chains (width of yellow dots in Fig. 4a, b, and f). Additionally, it slightly increased the separation of synfire chains (Supplementary Fig. 4e). The shorter synfire chains led to a lower probability of correctly predicting reward (Fig. 4g, h), thus resulting in a decreased prediction rate (Fig. 4i), and the temporal extension of self-location increased the likelihood of avoiding traps and leading to a slightly higher dodge rate (Fig. 4j). Nevertheless, the substantially reduced prediction rate resulted in a worse overall rate of reward acquisition (Fig. 4k). Our findings suggest that gamma oscillations played a beneficial role in generating longer synfire chains, consequently enhancing predictive capabilities.

### 2.7 Trade-off between Theta Frequency and Motion Velocity Optimizing Expected Reward

The positive correlation between theta frequency (*f*_*θ*_) and motion velocity (*v*) can now be examined by exploring their functional interplay in optimizing the goal-navigation performance. Here, synfire chain dynamics were employed to investigate the influence of these factors on the prediction and dodge rates during navigation. Our simulations revealed that increasing *f*_*θ*_ consistently shortened synfire chains, while *v* had a relatively weaker impact on synfire chain length (Fig. 5a). Conversely, elevating *f*_*θ*_ decreased synfire chain separation, whereas increasing *v* augmented the separation (Fig. 5b). Shorter synfire chains resulted in reduced prediction distances, limiting the ability to reach the reward site and consequently lowering prediction rates at higher *f*_*θ*_ (Fig. 5c). On the other hand, although *v* weakly influenced synfire chain length, higher *v* reduced the number of synfire chains executed before reaching the fork (Supplementary Fig. 5) and diminished the predictive opportunity, thereby decreasing the prediction rate (Fig. 5c).

**Fig. 5.**
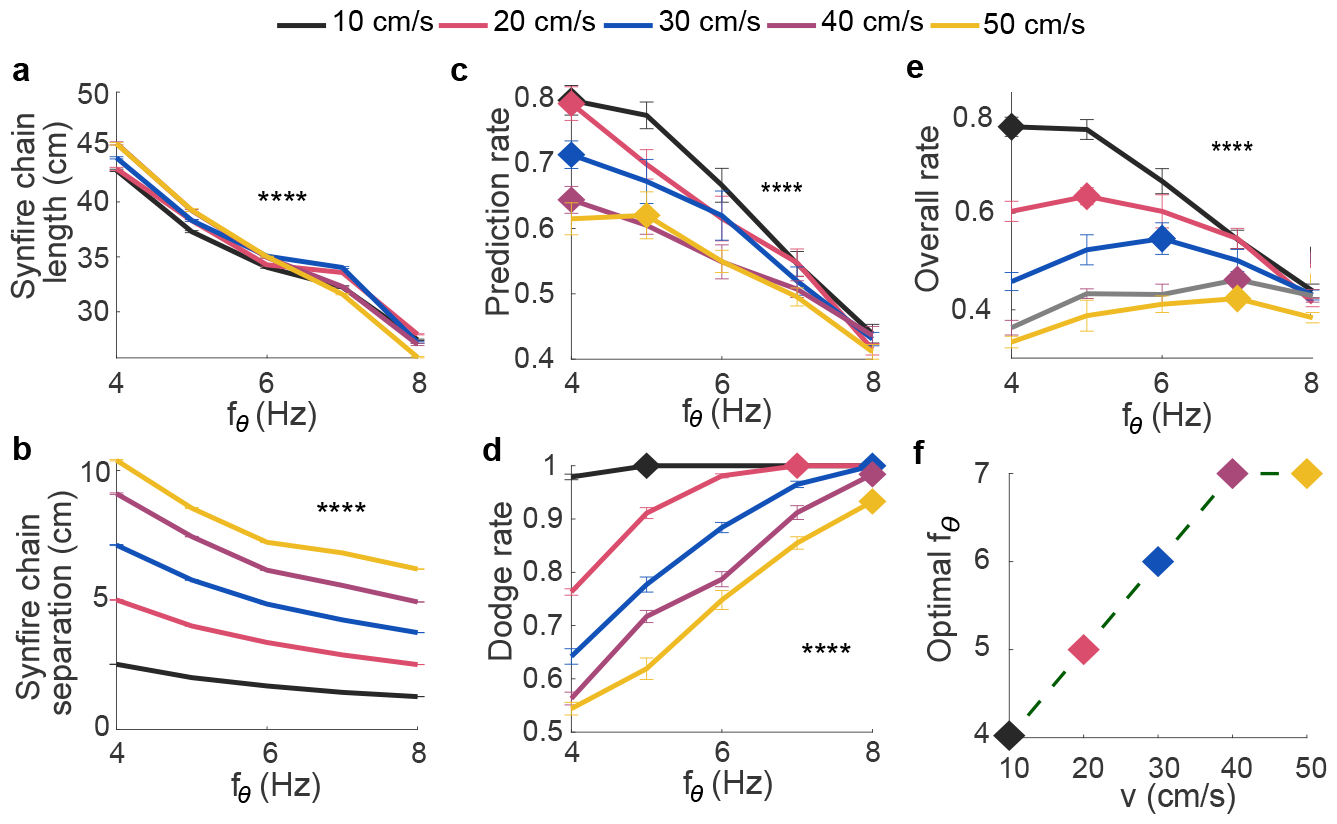
Optimal theta frequency at different motion velocities. (a, b) Synfire chain properties at various *v* and *f*_*θ*_: Synfire chain length (a); Synfire chain separation (b). (c-e) Goal-navigation performance: (c) Prediction rate; (d) Dodge rate; (e) The overall rate of reaching rewards. Diamond: maximum point on each curve at each velocity. (f) Dependence of optimal *f*_*θ*_ on *v*. Data in (a-e) are represented as mean *±* SEM. *n* = 10 trails/setting. ****: *p <* 0.0001, 2ANOVA test.

Furthermore, our findings indicated that increasing *f*_*θ*_ led to more extensive coverage of the main track by self-locations, enhancing the probability of successfully avoiding traps (Fig. 5d). Conversely, elevating *v* reduced this probability by increasing synfire chain separation (Fig. 5d). Considering the mean velocity *v* and consecutive synfire chains temporally separated by approximately the theta period (1*/f*_*θ*_), the expected spatial separation around *v/f*_*θ*_ was validated in our simulations (Fig. 5d). Thus, increasing *f*_*θ*_ decreased the temporal separation between self-locations of consecutive synfire chains, resulting in a higher dodge rate. Conversely, increasing *v* amplified synfire chain separation within each 1*/f*_*θ*_ interval, yielding a lower dodge rate (Fig. 5d). Overall, our observations consistently indicated that lower *f*_*θ*_ was more effective in predicting rewards (Fig. 5c) but less proficient in avoiding traps (Fig. 5d). However, with increasing *v*, both prediction and dodge rates decreased (Fig. 5c, d), albeit less significantly at higher frequencies. Consequently, the optimal overall rate gradually shifted towards higher *f*_*θ*_ as *v* increased (Fig. 5e, f). These findings underscore the trade-off between predicting future events and maintaining situational awareness of the current surroundings, elucidating the phenomenon of theta frequency increasing with motion velocity.

## 3 Discussion

The investigation conducted in this study sheds light on the intricate dynamics of synfire chains across varying theta frequencies and movement velocities, while also examining the consequences of impaired theta or gamma oscillations. Our findings underscore the significance of the positive correlation between theta frequency and movement velocity, revealing insights into the brain’s inherent capacity to anticipate future events (*synfire chain length*) and maintain a heightened awareness of surroundings (*synfire chain separation*).

Within this framework, impairment in theta oscillation significantly compromises both predictive capabilities and dodge ability, manifesting as a reduction in synfire chain chunking ability and a loss of self-location. Impairment of gamma oscillation primarily diminishes prediction ability due to shortened synfire chain length. These observations highlight the critical roles of theta and gamma oscillations in facilitating effective navigation and decision-making.

Furthermore, our study unveils a nuanced relationship between movement velocity and theta frequency. As subjects move faster, the rapidly changing environment necessitates increased vigilance and responsiveness to sudden changes. However, heightened alertness comes at the cost of reduced available time for predicting future events, resulting in a diminished range over which predictions can be made. Consequently, slower velocities favor lower theta frequencies, allowing more time for prediction, while higher velocities favor higher theta frequencies, facilitating enhanced alertness and responsiveness to the rapidly changing environment. Therefore, the optimal theta frequency increases proportionally with motion velocity. This nuanced interplay between movement velocity and theta frequency underscores the adaptive nature of the brain’s navigation strategies.

### 3.1 Goal Navigation and Unexpected events

The complexity of goal navigation, intertwined with the selection of the optimal path towards a reward, is intricately tied to the interplay of various neural mechanisms, including synfire chains, theta-nested gamma oscillations, and spatial information processing in decision-making. Prior studies [2–4, 13, 17] have significantly advanced our understanding of these mechanisms, particularly regarding spatial cognition and the utilization of cognitive maps, a concept introduced over seven decades ago [41].

Synfire chains, integral to goal navigation, associate closely with the concept of cognitive maps. The formation of cognitive maps was not considered in our study, as unexpected events can not be involved in the value system of the existing framework: the successor model and predictive map [42–44]. Instead, our model has focused on leveraging synfire chains to read these maps, which were stored within synaptic connections. While previous research has primarily highlighted the use of synfire chains in predicting future outcomes during navigation, our study brings attention to their response to unexpected events. Navigating through unpredictable scenarios is a common challenge in natural environments, underscoring the importance of understanding how synfire chains adapt to such circumstances. Our findings suggest that while synfire chains excel at predicting future events, they may struggle with unexpected disruptions. This highlights the necessity of integrating unpredictability into goal-navigation tasks to better understand the robustness of neural navigation strategies. Our study reveals that while synfire chains facilitate future prediction, they may be susceptible to unpredictable events. The observed correlation between motion velocity and theta frequency hints at a compensatory mechanism, illustrating the brain’s adaptability to unforeseen events to maintain effective navigation strategies. Thus, our study offers novel insights into the functional role of synfire chains in navigation.

Theta-nested gamma oscillations, intertwined with synfire chains during navigation, are believed to be pivotal for memory formation and retrieval [1, 2]. Gamma oscillations orchestrate activity synchronization across brain regions [10, 11], aiding memory retrieval [10, 11]. They align with the time constant of spike-timing-dependent plasticity (STDP) [45, 46], facilitating more effective synaptic modification, especially in the occurrence of spike-gamma coherence [16, 47]. These properties instantiate their importance in memory formation as well as the formation of cognitive maps, indicating the significance of gamma oscillations in encoding spatial memory. Our investigation, centered on synfire chains in CA1 but not concerning the formation of cognitive maps, unveils that gamma oscillations within theta-nested gamma oscillations effectively extend synfire chains, and thus enhance memory performance. This underscores the significance of gamma oscillations in CA1 for retrieving rewards via synfire chains. On the other hand, theta oscillations within theta-nested gamma oscillations serve to chunk spatial information during locomotion [7, 8]. They also play a crucial role in memory retrieval, evidenced by increased theta power during correct recall [4, 6]. Our study delves into spatial information retrieval through synfire chains, which were modulated by theta-nested gamma oscillations. Our findings suggest that theta oscillations not only chunk spatial information but also orchestrate synfire chains, whose disruption led to poorer learning between L1 and L2, subsequently impacting recall performance. This underscores theta oscillations’ significance in learning reward location during goal-navigation tasks.

Navigation also intricately intertwines spatial information processing with decision-making, primarily orchestrated by the prefrontal cortex (PFC) [48, 49], while the hippocampus, housing diverse cell types like place cells, head direction cells, and reward cells, crucially contributes to spatial representation and navigation [17, 49]. However, recent experimental findings propose an expanded role of the hippocampus, suggesting its involvement in working memory and decision-making processes. Activation of memory engrams in CA1 enhances memory retrieval [50, 51] and the hippocampus can store multiple instances of working memory through cross-frequency coupling [52], indicating its broader functional scope beyond spatial processing. Additionally, the activity of splitter cells in CA1 has been linked to predicting future decisions, further suggesting CA1’s involvement in working memory and decision-making processes [53–55].

Specifically, recent research highlights feedforward connections from L2 to PFC [28], indicating CA1’s involvement in providing reward-related cues for working memory and decision-making. During navigation, the hippocampus exhibits VTE activity at decision points[3, 18], where synfire chains activate and sweep through possible trajectories, and thus anticipate future actions based on spatial cues. Both CA3 and CA1 contribute to this process, integrating spatial information into decision-making. Retrieval of spatial information crucial for decision-making involves sensory inputs processed in the neocortex, routed through grid cells within EC before reaching the hippocampus, triggering synfire chains during VTE to anticipate forthcoming actions. Our model incorporates this by retrieving spatial information regarding self-location within a maze via location-dependent inputs, which navigate through grid cells to the hippocampus (Fig. 1a and 2b). Upon the synfire chain in L1 reaching the reward site, the neurons encoding the reward in L2 are long-lasting activated as engram cells, and subsequently transferred to PFC to facilitate working memory and decision-making. Activation of neurons encoding rewards in L2 positively correlates with goal-navigation performance, indicating their involvement in decision-making. Our study supports the notion that working memory and decision-making processes also occur in CA1, with the synfire chain length constraining the retrievable memory region.

### 3.2 Biological Plausibility of Our Model

Synfire chains and theta-nested gamma oscillations have been frequently modelled independently to elucidate their respective functions, yet their intricate interplay is not fully understood [47, 56, 57]. In light of recent experiments [1–3, 13, 17, 18, 35] supporting the interplay between goal navigation, synfire chains, and theta-nested gamma oscillations, we developed a biologically plausible computational model to investigate the functional roles of synfire chains under the modulation of theta-nested gamma oscillations in various navigational scenarios with both predictable and unpredictable events. Our model successfully reproduces several key phenomena observed in experiments, including theta-phase-gamma-amplitude cross-frequency coupling and theta-phase precession, when coupled with external theta input. Both theoretical considerations and experimental data indicate that theta-phase precession and theta sequences may underlie cognitive functions [1–3, 17].

Recent evidence has underscored the critical role of place cells in forming cognitive maps and generating synfire chains for navigation, emphasizing their unique features such as sparse and distance-dependent connectivity and distinct place fields [2, 17, 58]. As spatial information is stored in synaptic connections, aligning with the synaptic theory of memory storage [50, 51, 59], synfire chains were generated at the network level, consistent with the framework known as the continuous attractor model [2, 15, 60, 61]. In this framework, place cells are activated at corresponding place fields through location-dependent inputs to a circuit with distance-dependent connectivity and adaptation, leading to synfire chain propagation. Here, we employ this framework to simulate place cell activity by representing place fields as Gaussian functions [16, 47, 62–66], generating synfire chains triggered by location cues and decision points. Our model, incorporating symmetric connections and short-term depression [47, 56], aligns more closely with biological plausibility by temporally depressing synaptic conductance after the subject’s traversal, facilitating synfire chain generation aligned with theta sequences and movement direction.

Under this framework, the widely observed theta-nested gamma oscillations [3, 5, 39] can be naturally generated during navigation. Theta oscillation is believed to originate in MS [26, 67, 68], as its perturbation results in reduced theta power [4, 69, 70] and the two mutually inhibited populations MS1 and MS2 fire out of phase [21–23, 71]. This mechanism also facilitates the generation of theta-phase precession through modulation of location-dependent inputs [2, 15, 60]. Gamma oscillation, generated by the E-I loop within CA1 in our model, requires a time delay between excitatory and inhibitory spikes, which is supported by the experimental evidence: E-I phase delay in CA1 [33, 72, 73]. Thus, coupling MS and CA1 as suggested in our model is competent to generate robust theta-nested gamma oscillations during navigation, offering flexible control of theta and gamma oscillations independently.

Our model offers valuable insights into the mechanisms underlying theta-nested gamma oscillations, theta-phase precession and synfire chain generation [47, 56]. However, alternative models such as the Somato-Dendritic Model and the Oscillatory Interference Model provide different perspectives and contribute to a more comprehensive understanding of these phenomena [74–80]. Importantly, the various models proposed are likely to yield consistent results as long as they produce synfire chains with theta-nested gamma oscillations and theta-phase precession, ensuring robustness and reliability across different modelling approaches. Although challenges persist in fully integrating all components into a unified computational model, our study marks a significant advancement in elucidating their functional roles in navigation.

### 3.3 Limitation of the Study

Our model is simplified by excluding spike-timing-dependent plasticity (STDP) within L1. This decision overlooks potential influences of theta frequency and motion velocity on synfire chain learning, which could alter spike-time pairings. Further investigations are warranted to explore these effects in greater detail.

While the positive correlation between theta frequency and motion velocity has long been acknowledged, it remains understudied despite advancements in models concerning synfire chains, theta oscillations, and learning [13, 14, 81]. Our study proposes that a trade-off between synfire chain length and separation may underlie the necessity for this correlation. However, we acknowledge limitations in reproducing the phenomenon of synfire chain slope increasing with movement velocity, which may affect the proposed trade-off, as increasing the slope potentially extends prediction distance at higher velocities.

To address this discrepancy, plasticity should be introduced to elevate excitatory input or reduce inhibitory input to CA1 neurons [82–84]. A recent study suggested that Hebbian plasticity applied to synaptic weights during the self-location phase of a synfire chain could potentially generate the desired increase in synfire chain slope with movement velocity [82]. The increased slope of synfire chains would further improve the subject’s overall performance, providing better prediction ability without affecting, as presented in Fig. 2i, where optimal ones were located at the low right corner. Thus, this finding provides valuable insight for further exploration into the intricacies of the effect.

Additionally, our exploration could be expanded by introducing more complex goal-navigation tasks, such as incorporating multiple rewards at varying distances from a fork, to scrutinize dependence on reward locations relative to each other. While our study proposes a trade-off between theta frequency and motion velocity, suggesting an optimization in reward acquisition, the absence of maze studies with trap settings necessitates future animal experiments to validate our findings.

In summary, our study offers significant insights into the intricate interplay among synfire chains, theta-nested gamma oscillations, movement velocity, and navigation abilities, deepening our understanding of the neural mechanisms underlying goal navigation and laying the groundwork for more effective strategies in navigating complex environments. Looking forward, further research could explore interventions or training protocols to enhance navigation skills across diverse conditions, advancing neuroscience and practical applications like robotics and human-machine interaction. Additionally, our investigation sheds new light on the functional role of synfire chains in navigation, emphasizing the importance of adaptability in neural navigation models. By unravelling the complexities of goal navigation and spatial cognition, we edge closer to creating resilient navigation systems, offering potential for innovative approaches to navigating intricate environments.

## 4 Methods

### Circuit architecture and dynamics

Our model featured a tri-layered structure, comprising MS, L1 and L2. MS comprises 500 inhibitory neurons, divided into MS1 and MS2, each consisting of 250 inhibitory neurons [21, 22]. Neurons were randomly connected with a probability of *p* = 0.2, where synaptic conductance 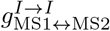 was set for mutual inhibition between MS1 and MS2, and 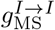 for self-inhibition insider each subpopulation. Both L1 and L2 consisted of 2000 excitatory neurons (*n*_*E*_), and 500 inhibitory neurons (*n*_*I*_). Each excitatory neuron *i* was associated with a preferred place field *x*_*i*_. Neurons within the same layer were randomly connected at a probability of *p* = 0.2, except for excitatory neurons, which were connected at a probability of 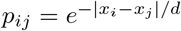, where *d* was the connectivity length. A periodic boundary condition was applied to the maze, with *x*_*i*_ *x*_*j*_ = 1 if neuron *i* and *j* encoded the starting point and end point of the maze respectively, resulting in a continuous attractor [85, 86].

As justified in the context of previous studies [36, 40, 71, 73, 84, 87], our biologically plausible model utilized conductance-based integrate-and-fire neurons, incorporating short-term depression and long-term plasticity mechanisms, with the dynamics of *i*th neuron’s membrane potential 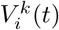 given as:

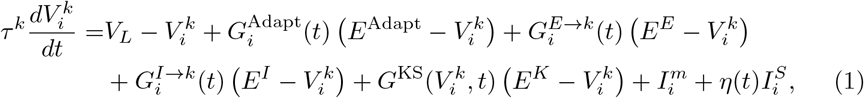

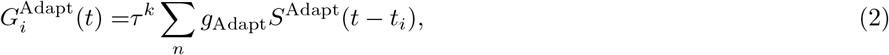

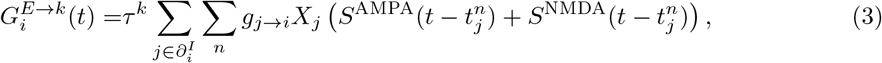

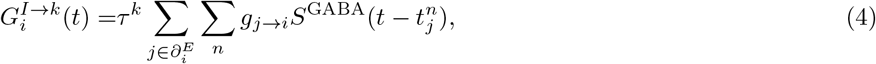

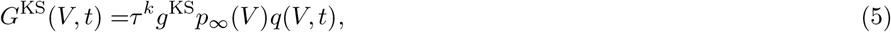

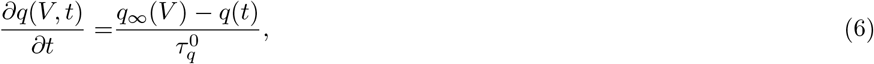

with *q*_∞_(*V*) = 1*/*(1 + *e*^(*V* +65)*/*6.6^) and *p*_∞_(*V*) = 1*/*(1 + *e*^−(*V* +34)*/*6.5^); *k* = *E* (*I*) signified excitatory (inhibitory) neurons and 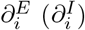denoted excitatory (inhibitory) presynaptic neurons of neuron *i*; 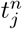 was the *n*th spike time of neuron *j*; *g*_*j*→*i*_ represented the synaptic strength from neuron *j* to neuron *i*, with *i, j* ∈ *E* or *I*; *g*_Adapt_ denoted the synaptic strength of adaptation only on excitatory neurons in L1, while *g*^KS^ denoted the conductance of slow potassium current only for neurons in MS; *S*^*α*^(*t*) represented the synaptic time course of various receptors *α* = AMPA, NMDA, GABA, modeled by a bi-exponential function with characteristic decay 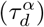 and rise time constants 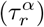, taking the form:

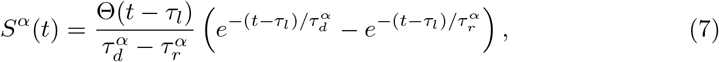

where Θ(*t*) was a Heaviside step function; *S*^Adapt^(*t*) denoted the time course of adaptation, taking a similar form with Eq. 7; Short-term depression [87, 88] was applied to excitatory synapses, described by the amount of neurotransmitter *X*_*i*_, whose dynamics were given as

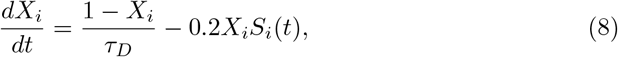

with a decay time constant *τ*_*D*_ and the spike train of neuron *i* given as 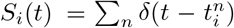, where *δ*(*t*) was the Dirac delta function. When the inter-spike interval was smaller than *τ*_*D*_, frequent depletion of neurotransmitters induced short-term depression, temporarily reducing synaptic efficacy.

Besides, long-term plasticity was applied to excitatory-to-excitatory (*E* → *E*) from L1 to L2, where R-STDP [37, 89] was employed, updating the synaptic efficacy *g*_*j*→*i*_ as follows:

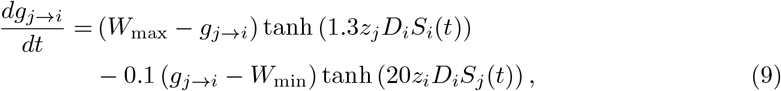

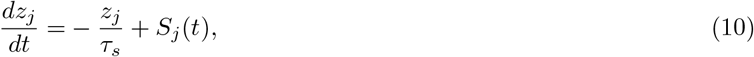

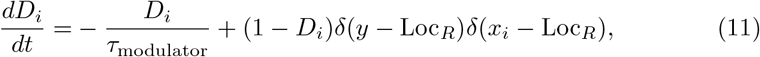

with *y* being the subject’s current location, *x*_*i*_ neuron *i*’s preferred place field and Loc_*R*_ the reward location; *z*_*j*_ was the synaptic trace induced by spikes from neuron *j*, while *D*_*i*_ was the neuromodulator signal at neuron *i*, tagging neuron *i*’s pre-synaptic efficacies modifiable; Thus, only those *g*_*j*→*i*_ with highly activated neurons and tagged post-synaptic neurons can undergo such R-STDP.

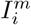 denoted the external input, including the background input *I*_MS_ from MS and the location-dependent input *I*_L_ received by excitatory neurons, as introduced in Section 2.1, taking the form:

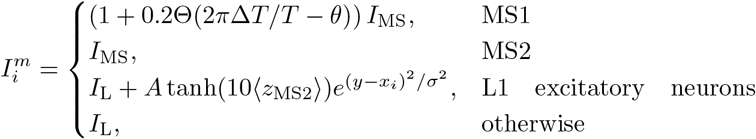

where Θ(Δ*θ*) was a Heaviside function, with Δ*T* denoting the off-period of time within the theta period *T*, as shown in Fig. 1a, and *θ* denoting the phase of theta oscillations; *y* was the subject’s current location, while *x*_*i*_ was the preferred place field of neuron *i*; ⟨*z*_MS2_⟩ was the mean synaptic trace of the population MS2; *A* and *σ* were the amplitude and standard deviation of the CA3 input, respectively. 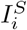 represented the amplitude of background fluctuation and *η*(*t*) denoted a random number generator in the range [− 0.5, 0.5).

All the above parameters in our model were summarized in Tab. A1.

### Simulation details

The linear maze depicted in Fig. 1b consisted of 2000 excitatory neurons. It spanned 200 distinct locations, with each location Loc_*I*_ spanning 1 cm and encoded by 10 neurons, e.g., Count(*x*_*i*_∈ Loc_*I*_) = 10. The cross maze illustrated in Fig. 2a was divided into four sections with each section covering 50 locations and represented by 500 excitatory neurons in both L1 and L2. In the learning phase, the rat started from the main track and sequentially visited each arm at least twice.

Upon reaching the randomly assigned reward site, a neuromodulator signal was induced to the excitatory neurons in L2, which encoded the reward site (*x*_*i*_ = Loc_*R*_), thus activating R-STDP to strengthen the excitatory synapse at the reward site (Supplementary Fig. 1b, d). The task phase commenced after the learning phase, and R-STDP was no longer applicable. During this phase, the rat moves at a variable velocity *v*∈ [0, 2*v*_0_] with *v*_0_ being its mean.

Upon reaching the fork of the cross maze, the rat faced a decision point, choosing arm *i* at the probability 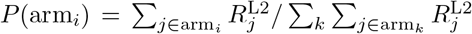 where 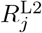 denotes the firing rate of neuron *j* in L2. The trap was randomly assigned to occupy 3 cm length along the main track. When the rat got close to the trap region, each location Loc_*I*_ ‘s total synaptic trace was calculated every ms, that is, 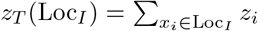, and 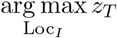within the trap region indicated that the rat had recognized and successfully dodged the trap, continuing to seek the reward. Otherwise, it was trapped and the trial terminated.

### Synfire chain detection

To detect synfire chains, we first identified the *a*th time 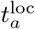 when place cells of the subject’s current location were activated (yellow dots in Fig. 1d), that is, the coincidence of *y* and 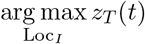. A synfire chain was defined as spike trains fired during the period 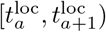 for any *a*. The displacement during this period defined synfire chain length, formulated as 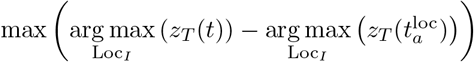. Synfire chain separation was defined as the spatial 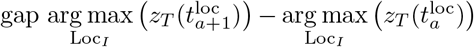. The number of synfire chains was counted from the starting point of the cross maze to the fork along the main track.

### Measurements

Oscillation power in L1 was computed by Fast Fourier Transforming (FFT) the detrended mean membrane potential of all excitatory neurons in L1 within a 10-second interval, with the summation of power in the theta band [4, 8] Hz for theta power, and that in the gamma band [25, 40] Hz for gamma power. Subsequently, an inverse FFT was applied to obtain the band-passed theta and gamma oscillations, respectively.

The intensity of theta-gamma phase-amplitude coupling can be measured by a modulation index (MI) as described [9]. MI can be obtained by calculating the Kullback-Leibler (KL) divergence *D*_KL_(*D* || *U*), where *D* represents a normalized distribution of the mean gamma amplitude across various theta phases binned into 18 20^*°*^ intervals (0 ∼ 360^*°*^), while *U* is a uniform distribution over theta phases. A higher MI signifies a stronger coupling between theta phase and gamma amplitude.

Neuronal synchrony *K*_*ij*_ measures the probability of the neuron pair *i* and *j* firing in coincidence, given as 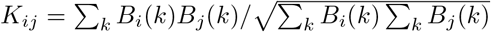, with *B*_*i*_(*k*) (*B*_*j*_(*k*)) being the spike train of neuron *i* (*j*). *B*_*i*_(*k*) = 0 (1) denotes no spike (one spike) generated within the *k*th 1-ms bin. The average of all pairs of excitatory neurons in L1 was defined as the synchrony index.

The field information [4] was employed to quantify the correlation between the subject’s location and the synfire chains in L1, that is, how well neuronal activities encoded the spatial information. It was defined as 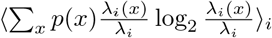 denotes the location in the maze, and *p*(*x*) is the probability of the subject occupying the location *x*; *λ*_*i*_(*x*) represents the firing rate of neuron *i* at location *x*, while *λ*_*i*_ denotes the average firing rate of neuron *i* across the entire maze. A higher value of the field information indicates a better encoding of the subject’s location by the activity of excitatory neurons in L1.

Theta-phase precession was quantified by the circular-linear correlation (clcorr) between the theta phase and the distance to the neuron’s preferred location. It was calculated here using the toolbox [90], based on the formula 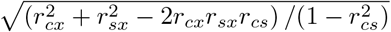, where *r*_*cx*_ = corr(cos(*θ*_theta_), *x*), *r*_*sx*_ = corr(sin(*θ*_theta_), *x*) and *r*_*cs*_ = corr(cos(*θ*_theta_), sin(*θ*_theta_)), with *x* denoting the distance between spiking location and preferred location, corr and *θ*_theta_ being the Pearson correlation and the theta phase, respectively. A higher clcorr indicates that the spike locations away from the preferred location correlate more with the theta phase.

## 5 Data availability

The data that support the plots within this paper and other findings of this study are available from the corresponding author upon reasonable request.

## Acknowledgements

This work was supported partly by the National Natural Science Foundation of China (Grant No. 12175242), the Natural Science Foundation of Zhejiang Province (Grant No. LZ24A050007), the Research Initiation Project of Zhejiang Lab (No. K2022KI0PI01) and the Zhejiang Lab Open Research Project (No. K2023KI0AA03).

## Author contributions

K.T.L., Y.W., P.G. and D.Y. conceived the study. K.T.L. performed the model simulations and data analysis. K.T.L., Y.W., P.G. and D.Y. participated in discussion and wrote the paper.

## Additional information

Supplementary material in Appendix A.

### Appendix A Supplemental Figure and Table

**Fig. A1.**
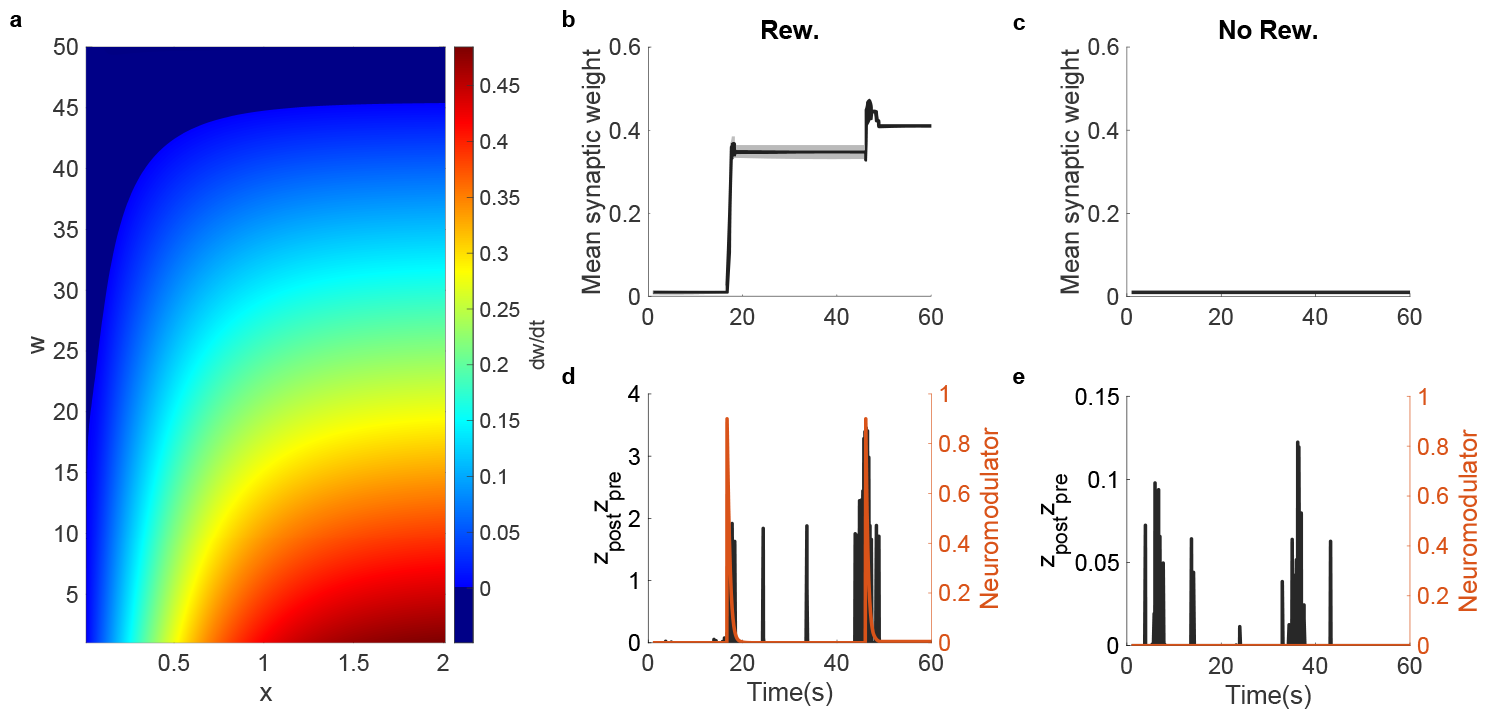
(Related to Fig. 2) Synaptic weight evolution. (a) Synaptic weight change through replacing *z*_*j*_ *D*_*i*_*S*_*i*_(*t*) and *z*_*i*_*D*_*i*_*S*_*j*_ (*t*) by *x* in Eq. 9. (b, c) L1 to L2 synaptic weight at the reward site (the middle of one of the arms) (b) and not at the reward site (another arm of the maze) (c). (d, e) Coupled synaptic trace, *z*_pre_ and *z*_post_ are synaptic traces of pre- and post-synaptic neurons respectively, and post-synaptic neuromodulators at the reward site (d) and non-reward site (e). Data in (b, c) are represented as mean *±* standard error (SEM).

**Fig. A2.**
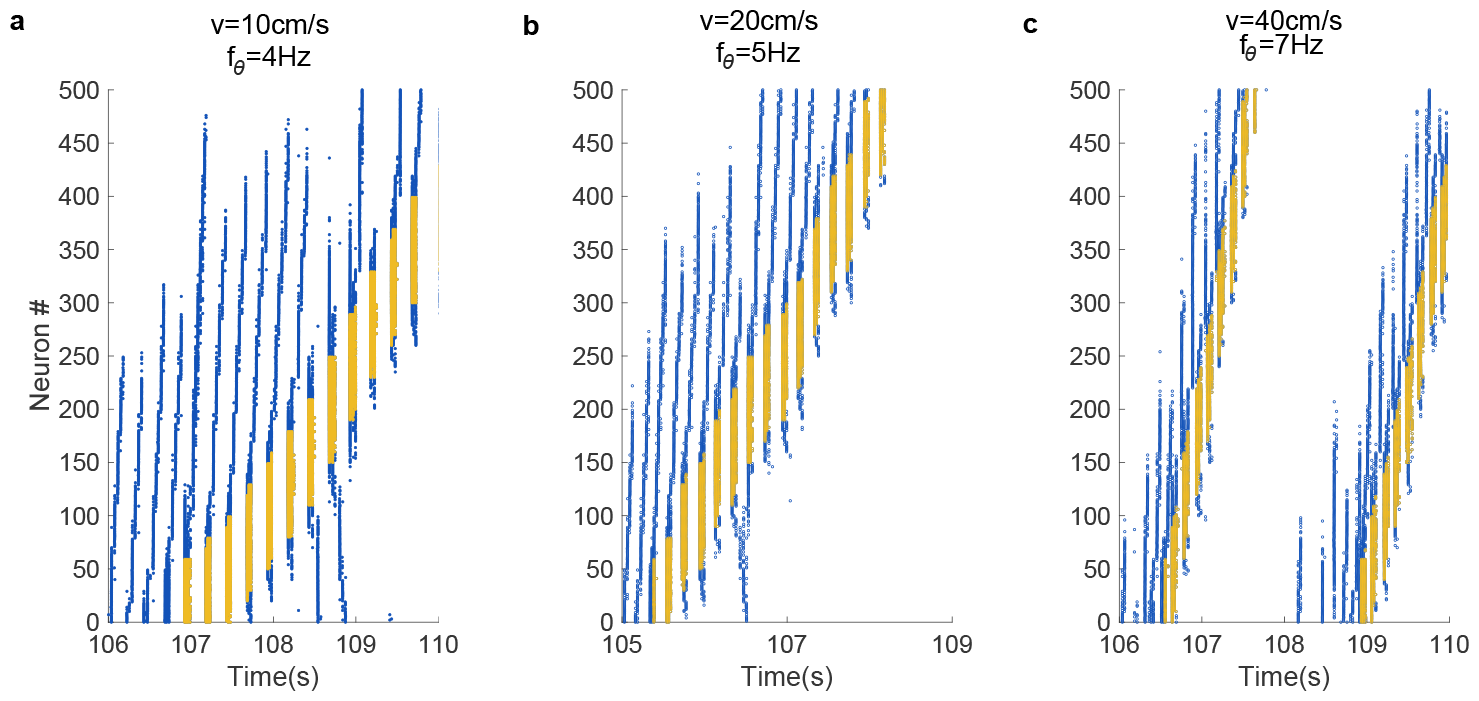
(Related to Fig. 2) Spike raster plots of synfire chains at various *v* and *f*_*θ*_. (a) *v* = 10 cm/s, *f*_*θ*_ = 4 Hz; (b) *v* = 20 cm/s, *f*_*θ*_ = 5 Hz; (c) *v* = 40 cm/s, *f*_*θ*_ = 7 Hz.

**Fig. A3.**
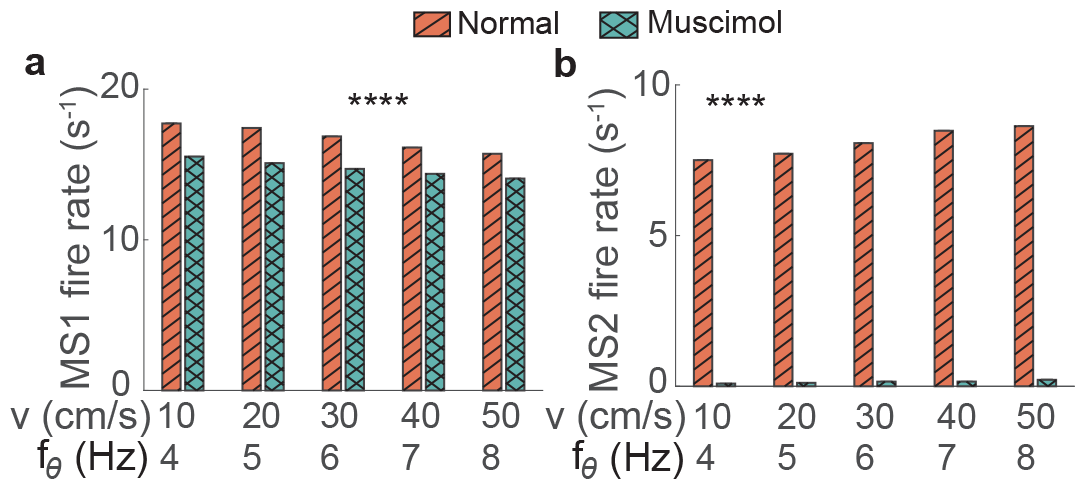
(Related to Fig. 3) Muscimol-induced firing rate reduction. Firing rate of MS neurons of MS1 (a) and MS2 (b). Data are represented as mean *±* SEM. *n* = 10 trails/setting. ****: *p <* 0.0001. 2ANOVA test.

**Fig. A4.**
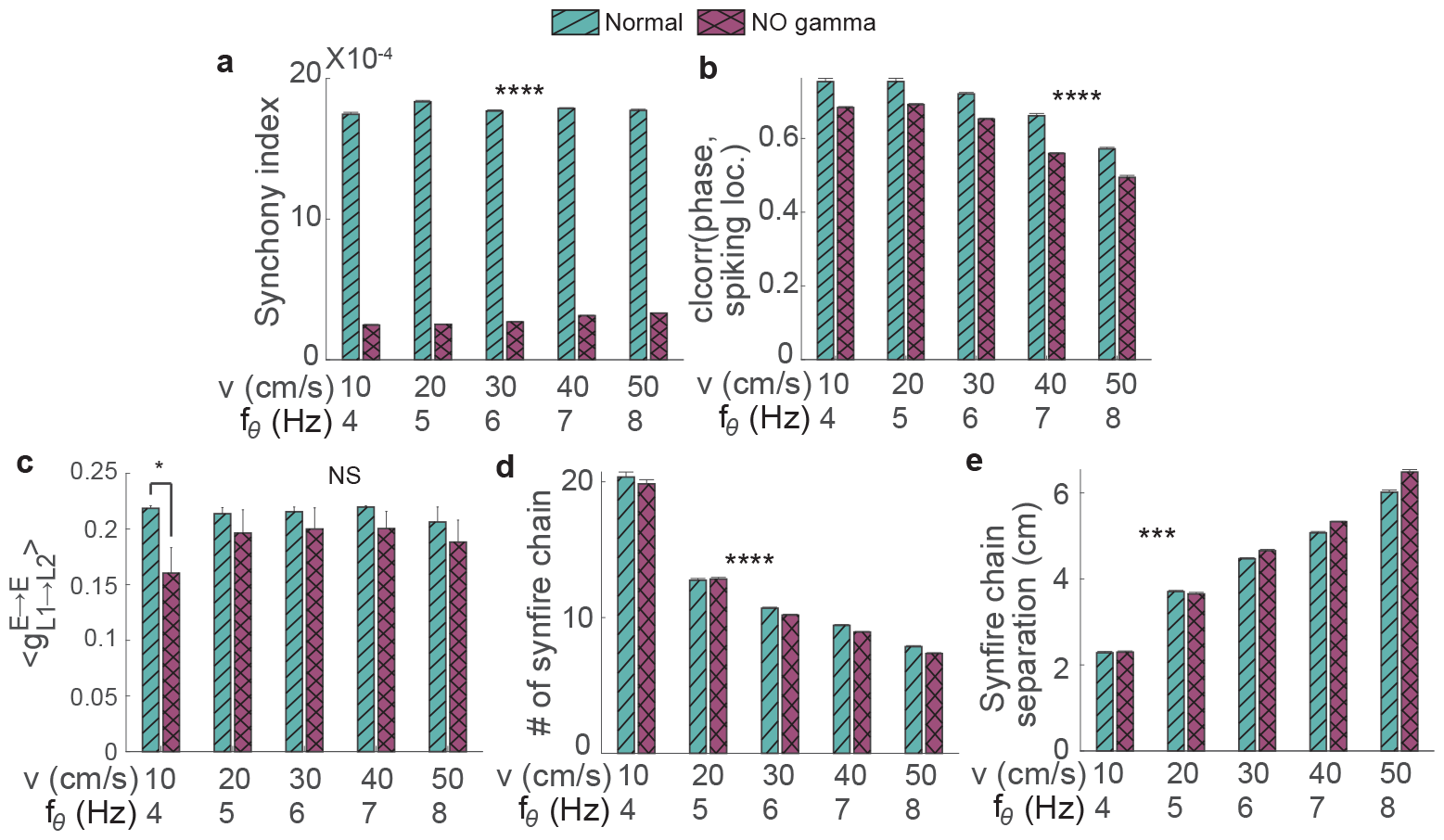
(Related to Fig. 4) Gamma oscillation impairment induced changes in synfire chain properties. (a) Synchrony index. (b) Circular-linear correlation (clcorr) between theta phase and spiking location. (c) The average excitatory to excitatory synaptic weight from L1 to L2. (d) number of synfire chains between the starting point and the fork. (e) Synfire chain separation. Data are represented as mean *±* SEM. *n* = 10 trails/setting. NS, *p >* 0.05; ***, *p <* 0.001; ****; *p <* 0.0001. *v* = 10 cm/s in (c), Two-sample Student’s t-test.

**Fig. A5.**
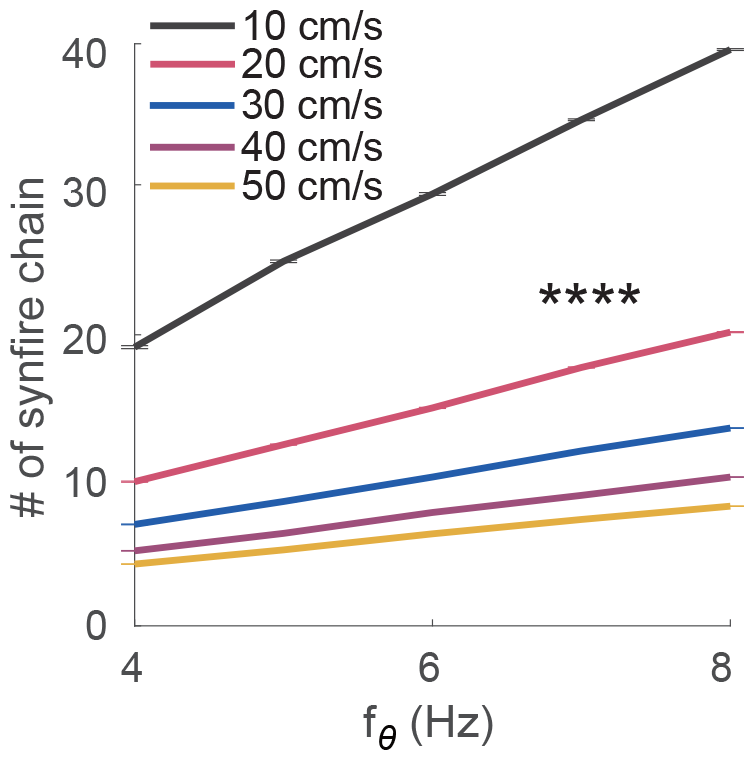
(Related to Fig. 5) Number of synfire chains from the starting point to the fork at various theta frequencies (*f*_*θ*_) and movement velocities (*v*). Data are represented as mean *±* SEM. *n* = 10 trails/setting. ****: *p <* 0.0001. 2ANOVA test.

**Table A1.** Simulation parameter summary

| Populations |  |  |
| --- | --- | --- |
| Description | Symbol | Value |
| Numbers of E, I neurons in L1 and L2 | $n_E, n_I$ | 2000, 500 |
| Numbers of I neurons in MS1 and MS2 | $n_{MS1}, n_{MS2}$ | 250, 250 |
| Connectivity in MS | $p$ | 0.2 |
| Connectivity length within L1 and L2 | $d$ | 5 cm |

| Neuron Model |  |  |
| --- | --- | --- |
| Description | Symbol | Value |
| Background input strength | $I_L, I_{MS}$ | 8 mV (L1, L2),<br>35 mV (MS) |
| Amplitude and standard deviation of location-dependent input | $A, \sigma$ | 150 mV, 5 mV |
| Axonal delay | $\tau_l$ | 1 ms |
| Leakage potential | $V_L$ | −60 mV (L1, L2), −40mV (MS) |
| Threshold potential | $V_{th}$ | −50 mV |
| Resting potential | $V_{rest}$ | −70 mV (L1, L2), −85 mV (MS) |
| Reset potential | $V_{rest}$ | −70 mV (L1, L2), −85 mV (MS) |
| Membrane time constant of E, I neurons | $\tau^E, \tau^I$ | 20 ms, 10 ms |
| Refractory period for E, I neurons | $t_{ref}^E, t_{ref}^I$ | 2 ms, 1 ms |
| Reversal potential of E, I neurons, potassium channel and adaptation | $E^E, E^I, E^K, E^{Adapt}$ | 0 mV, −85 mV, −85 mV, −85 mV |
| Leakage conductance | $g_L$ | 10 nS |
| conductance from E to E, I to E, E to I, I to I in L1 | $g_{L1}^{E \rightarrow E}, g_{L1}^{I \rightarrow E}, g_{L1}^{E \rightarrow I}, g_{L1}^{I \rightarrow I}$ | 0.1, 0.14, 0.8, 0.24 (normalized by $g_L$) |
| conductance from E to E, I to E, E to I, I to I in L2 | $g_{L2}^{E \rightarrow E}, g_{L2}^{I \rightarrow E}, g_{L2}^{E \rightarrow I}, g_{L2}^{I \rightarrow I}$ | 0.04, 0.3, 0.28, 0.48 (normalized by $g_L$) |
| Initial conductance from E in L1 to E in L2 | $g_{L1 \rightarrow L2}^{E \rightarrow E}$ | 0.01 (normalized by $g_L$) |
| Conductance between MS1 and MS2 and their self-inhibition | $g_{MS1 \leftrightarrow MS2}^{I \rightarrow I}, g_{MS}^{I \rightarrow I}$ | 0.03, 0.01 (normalized by $g_L$) |
| Conductance from MS1 to I in L1 | $g_{MS1 \rightarrow I}^{I \rightarrow I}$ | 0.375 (normalized by $g_L$) |
| Decay time constant of AMPA, GABA, and NMDA current | $\tau_d^{AMPA}, \tau_d^{GABA}, \tau_d^{NMDA}$ | 5 ms, 5 ms, 50 ms |
| Rising time constant of AMPA, GABA, NMDA current | $\tau_r^{AMPA}, \tau_r^{GABA}, \tau_r^{NMDA}$ | 0.5 ms, 0.5 ms, 0.5 ms |
| Adaptation strength | $g^{Adapt}$ | 10.5 (normalized by $g_L$) |
| Decay and rising time constant of adaptation | $\tau_d^{Adapt}, \tau_r^{Adapt}$ | 50 ms, 0.5 ms |
| Conductance of slow potassium current | $g^{KS}$ | 0.315 (normalized by $g_L$) |
| Time constant of slow potassium channel | $\tau_q^0$ | 189 ms |

| Short-term Depression |  |  |
| --- | --- | --- |
| Description | Symbol | Value |
| Time constant of short-term depression | $\tau_D$ | 200 ms |

| Reward Spike-timing-dependent Plasticity |  |  |
| --- | --- | --- |
| Description | Symbol | Value |
| Upper and lower limit of synaptic weight | $W_{\max}, W_{\min}$ | 0.5, 0.01<br>(normalized by $g_L$) |
| Time constant of spike-timing-dependent plasticity | $\tau_s$ | 20 ms |
| Time constant of neuromodulatory signal | $\tau_{\text{modulator}}$ | 1000 ms |

